# Broadleaved hedgerows as complementary habitats for small mammals in pine plantation landscapes

**DOI:** 10.64898/2026.03.17.712293

**Authors:** Aloïs Berard, Nattan Plat, Julien Pradel, Maxime Galan, Anne Loiseau, Sylvain Piry, Julie Blanchet, Lily Cesari, Karine Berthier, Jean-Baptiste Rivoal, Pellett Cameron, Valbuena Ruben, Hervé Jactel, Nathalie Charbonnel

## Abstract

1. The global decline of natural forests is accompanied by a rapid expansion of commercial tree plantations, which are expected to further increase to meet growing demand for wood products. However, planted forests generally support lower biodiversity than natural forests, particularly when monospecific and intensively managed. In this context, broadleaved hedgerows have been proposed as a nature-based solution to enhance biodiversity within conifer-dominated plantation landscapes. Such features may be especially beneficial for small mammals, including rodents and shrews, which are key contributors to forest ecosystem functioning. However, their effects on small mammal communities remain largely unquantified.
2. Here, we assessed variation in small mammal communities among habitat types within a native pine plantation-dominated landscape in southwestern France. Using a multi-year, multi-season survey, we compared species richness and abundance among plantation edges, broadleaved hedgerows embedded within plantations and natural broadleaved forests. We further tested whether environmental descriptors of hedgerow sites influenced dominant species and whether seasonal and interannual demographic dynamics modified habitat-related patterns.
3. Pine plantation edges and broadleaved hedgerows supported lower small mammal species richness than natural broadleaved forests and were dominated by two habitat generalists, *Apodemus sylvaticus* and *Crocidura russula*. This pattern was driven by the near absence of the forest specialist *Clethrionomys glareolus*. Hedgerows did not increase species richness relative to plantations, but provided favourable habitat for *A. sylvaticus,* which was scarce in pine plantation, while supporting fewer *C. russula*. Variation in hedgerow structure and composition further influenced *A. sylvaticus* abundance, while seasonal and interannual rodent population dynamics modulated habitat-related differences.
4. Our results indicate that intensively managed pine plantations act as environmental filters, excluding forest-associated small mammals. While broadleaved hedgerows benefited one species, their capacity to restore forest-specialist communities was limited without broader landscape-scale interventions. These findings highlight both the ecological benefits and constraints of edge-based habitat interventions and provide guidance for designing and evaluating biodiversity-oriented management in plantation landscapes.

## 1 INTRODUCTION

Forest ecosystems harbor the largest share of terrestrial biodiversity (Vié *et al*., 2009), yet this biodiversity and the ecosystem functions it supports (Brockerhoff *et al*., 2017) are increasingly threatened by human pressures (Green *et al*., 2020).

While the global extent of natural forests declines, the area covered by plantation forests, defined by the FAO as planted forests composed of even aged stands of one or two tree species, is increasing and is expected to expand further to meet rising production demands (FAO, 2025). Although plantations can provide alternative habitats to wildlife within humans modified landscapes, they generally support lower levels of biodiversity than natural forests owing to their compositional and structural homogeneity (Brockerhoff *et al*., 2008; Wang *et al*., 2022).

Multi-scale environmental heterogeneity, particularly vegetation heterogeneity, is a major driver of species diversity because it promotes species coexistence and persistence through niche differentiation, resource diversification and ecosystem stability (Stein *et al*, 2014; Heidrich *et al*., 2020). Beyond heterogeneity, biodiversity is also shaped by habitat amount. Under the Habitat Amount Hypothesis, species richness at a given location depends primarily on the total amount of habitat within the surrounding landscape (Fahrig, 2013). As plantations occupy large areas within landscapes, the total amount of natural forest habitat is expected to decline. A reduction in natural forest cover should therefore lead to reduced forest species richness, even where local patches of natural forest remain.

In plantation stands, both tree diversity and structural complexity are reduced. Even-aged management and low tree species richness reduce three-dimensional stand complexity (Juchheim *et al*., 2020) and are associated with lower understorey diversity (Kremer *et al*., 2025). Intensive management practices such as short rotation cycles, understorey clearing, thinning and deadwood removal further maintain stands in early to mid-successional stages and limit the development of structural features essential for forest-dwelling species (Tews *et al*., 2004; Wagenaar *et al*., 2025).

Monospecific plantations generally support lower plant and animal diversity than mixed-species ones (Wang *et al*., 2022; Kremer *et al*., 2025). Plant diversity influences animal diversity and abundance by driving the supply and variety of resources across trophic networks. Diverse plant communities increase the probability of fulfilling the trophic requirements of specialist herbivores (Siemann *et al*., 1998). Higher plant diversity may offer diet-mixing opportunities for generalists (Lefcheck *et al*., 2013) and increase overall resource availability by enhancing plant productivity (Chen *et al*, 2025), thereby supporting larger consumer populations and reducing extinction risk (Srivastava and Lawton, 1998). Increased richness and biomass of herbivores in turn sustains richer and denser predator communities through bottom-up trophic effects (Scherber *et al*., 2010). Beyond composition, structurally complex vegetation provides a greater diversity of microhabitats, shelter, and breeding sites, thereby expanding niche space and carrying capacity, and ultimately supporting higher animal diversity and abundance (Larrieu *et al*., 2022).

Enhancing animal diversity in plantation landscapes requires strategies that increase both environmental heterogeneity and total natural habitat amount. Current strategies focus on modification of stands composition (e.g. tree species diversification) and management practices (e.g. deadwood retention; Duflot *et al*., 2022; Muys *et al*., 2022), which may be difficult to implement without governance frameworks or incentives, particularly in landscapes dominated by small private landowners (Tiebel *et al*., 2022). A complementary approach consists in creating, restoring or preserving habitats at stands edges rather than modifying the stands themselves. These boundary habitats include features such as firebreaks or riparian strips in forests, and hedgerows and grassy margins in agricultural landscapes. Although they occupy a small proportion of total land cover, such semi-natural elements are widely recognised for their disproportionate value in enhancing landscape connectivity in human-modified environments (Arroyo-Rodríguez *et al*., 2020).

In agricultural settings, hedgerows (i.e. linear woody vegetation) provide habitat for many animal species and promote dispersal across poor quality matrices (Montgomery, Caruso and Reid, 2020). Transposing this approach to forest plantation systems has recently been conceptualised as a nature-based solution, with demonstrated benefits on pest regulation (Plat, Charbonnier, *et al*., 2025) and on the conservation of forest-dwelling species across multiple taxonomic groups (Plat *et al*., 2026). Nevertheless, small mammal responses to these features remain largely unaddressed despite their critical importance in forest ecosystem functioning and zoonotic transmission.

Rodents and shrews are core components of terrestrial food webs, particularly within forest ecosystems. Through seed predation and dispersal, fungal spore transportation and invertebrate consumption, they may influence plant regeneration, soil processes and community structures (Churchfield and Rychlik, 2006; Lacher *et al*., 2019; Borgmann-Winter *et al*., 2023). They also represent a major proportion of prey biomass for several predators, including raptors and carnivorous mammals (Jędrzejewska and Jędrzejewski, 1998). Beyond their ecological functions, small mammals are also of particular relevance for public health. They can host numerous zoonotic pathogens, including *Borrelia* spp., the causative agents of Lyme borreliosis (Radolf *et al*., 2012), and *Leptosira* spp. responsible for leptospirosis (Mayer-Scholl *et al*., 2014), among others. Small mammals also contribute to the maintenance of ectoparasite populations, including ticks and fleas, thereby playing a central role in vector-borne diseases dynamics (Ostfeld *et al*., 2018).

Furthermore, small mammals are highly sensitive to environmental conditions. Often relatively short-lived with limited home ranges, they can respond rapidly to environmental change on fine spatial and temporal scales (Andreassen *et al*., 2021). Tree species composition, silvicultural practices and landscape context shape interacting biotic and abiotic conditions to which small mammals respond in species-specific ways according to their ecological requirements (Fuentes-Montemayor *et al*., 2020). Consequently, small mammal assemblages exhibit varying degrees of habitat specialization, and their community composition can reflect underlying environmental conditions and disturbance regimes, thereby providing an indicator of ecosystem integrity (Avenant, 2011). Understanding the drivers of small mammal populations is also critical because fluctuations in their abundance can have cascading effects, including shifts in predator populations (Cano-Martínez *et al*., 2021) and altered pathogen circulation affecting zoonotic transmission (Khalil *et al*., 2019).

At the local scale, understorey structure and composition can influence the abundance of ground-dwelling small mammals by shaping resource availability and shelter from predators (Ecke *et al*., 2001). Abundances may also depend on broader drivers: the surrounding landscape can influence dispersal opportunities (Fitzgibbon, 1997), while climatic conditions such as rainfall can modulate demographic fluctuations through effects on vegetation productivity (Ventura-Rojas *et al*., 2025).

Consistent with these mechanisms, small mammal communities have been reported to exhibit reduced species richness and abundance in monospecific plantations relative to natural forests, with declines in forest specialists attributed to structural and compositional simplification (Happold and Happold, 1987; Chandrasekar-Rao and Sunquist, 1996; Pedersen *et al*., 2010). However, mammal responses to plantations remain poorly understood. Many studies have focused on exotic tree plantations, whereas native tree plantations may more closely resemble natural forests and thus support more diverse fauna (Stephens and Wagner, 2007). This limited and heterogeneous research cautions against broad generalisations and underscores the need for further research on small mammal responses.

In temperate regions, small mammal populations have been reported at lower abundances in conifer stands than in broadleaved ones, potentially reflecting reduced habitat complexity and reliance of some species on broadleaved-derived seeds (Fuller *et al*., 2004; Appleby and Balkenhol, 2024). Therefore, conifer plantations may combine structural and compositional constraints that negatively affect small mammal communities.

In this study, we assessed the potential of broadleaved hedgerows to enhance small mammal biodiversity within pine plantation-dominated landscapes. We conducted a multi-year, multi-season survey of small mammals in the Landes Forest (France), the largest intensively managed and human-planted forest in Europe. We examined how communities vary across habitat types. Specifically, we investigated whether (i) native pine plantations differ from natural broadleaved forests in terms of small mammal diversity and community composition; (ii) broadleaved hedgerows embedded within plantations can mitigate these differences; (iii) environmental descriptors of hedgerow sites influence dominant small mammal species; and (iv) temporal demographic dynamics influence these habitat-related patterns.

## 2 MATERIAL AND METHODS

### 2.1 Study sites and their environmental context

The study was conducted in the Landes de Gascogne forest in southwestern France, a 1.16 million hectare region dominated by native maritime pine (*Pinus pinaster*) plantations, which cover 66% of the area (Plat, Charbonnier, *et al*., 2025). These even-aged stands established on poor sandy podzol soils are intensively managed under rotation cycles of about 40-50 years, involving clear-cutting, soil preparation, thinning, and regular mechanical clearing of the understorey vegetation (Barbaro *et al*., 2005). Remaining land cover includes mostly agricultural land (15%), and unmanaged riparian forests and low-intensity managed broadleaved stands (13%), dominated by *Alnus glutinosa*, *Quercus robur*, and *Q. pyrenaica* (Plat, Charbonnier, *et al*., 2025). Within this pine plantation-dominated landscape, linear broadleaved structures occur along plantation edges, roads, forest tracks, and ditches. These semi-natural elements are maintained as property boundaries and for their cultural, recreational, and aesthetic value (Plat, Moreews, *et al*., 2025). We refer to them as “hedgerows”, by analogy with those found in agricultural landscapes.

In the Landes forest described above, our sampling was conducted across 53 sites distributed over a 1,050 km² area (center : 44°33’38.2”N 0°46’36.7”W), encompassing three habitat types : pine edges (12 sites), broadleaved hedgerows (24 sites) and broadleaved forests (17 sites; Figure 1). The sampling design followed Plat *et al*. (2026), with the addition of broadleaved forest sites located mainly in riparian zones and old forest remnants. Hedgerows were primarily composed of native oak species (*Q. robur* and *Q. pyrenaica*). They measured at least 100 meters in length, eight meters in height, and one to two trees in width, with continuous tree crowns.

**FIGURE 1.**
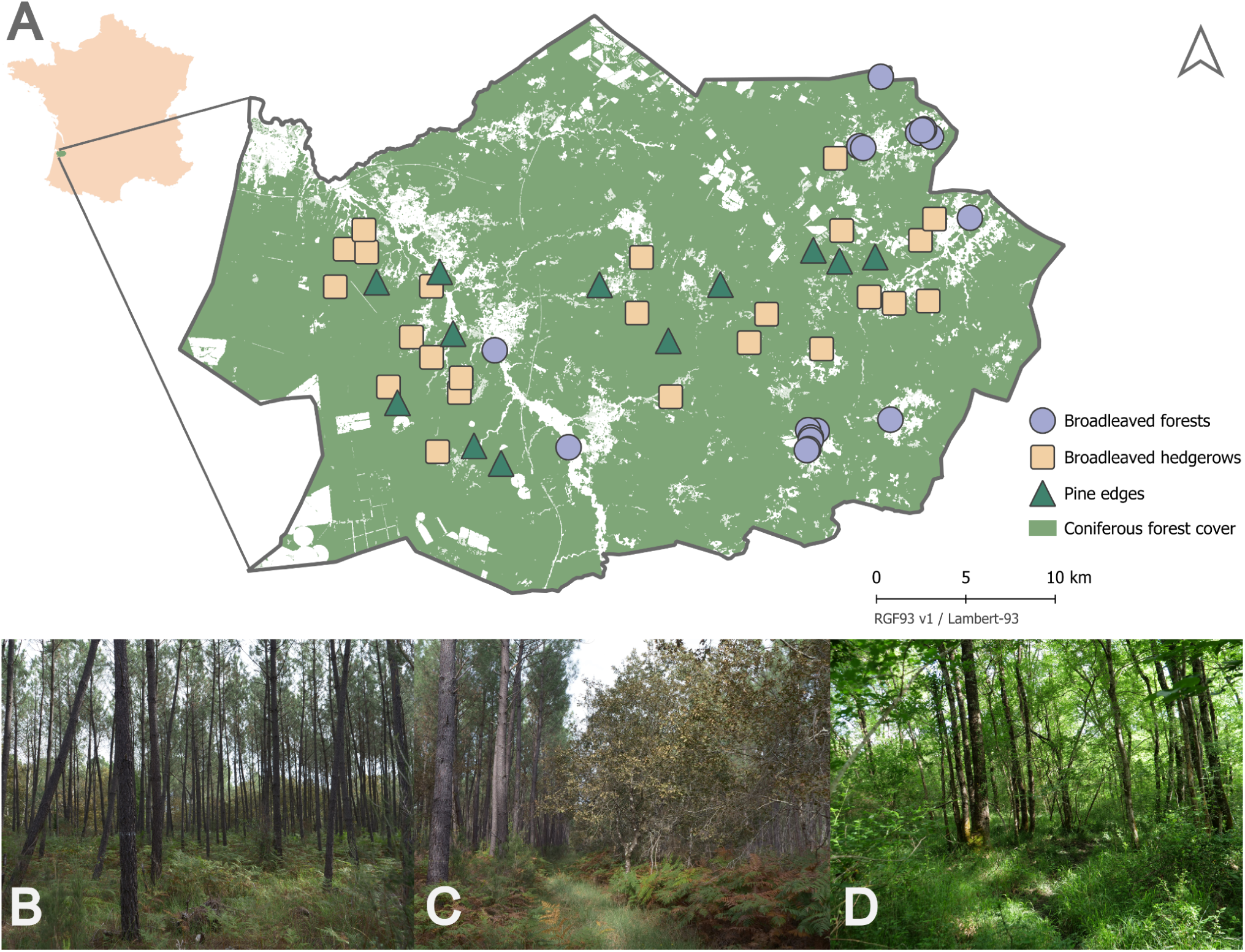
**A.** Map of sampling sites within the Landes forest (France). Coniferous forest cover data are produced by CNES for the Theia data centre (www.theia-land.fr) using Copernicus products. **B.** Pine plantation. **C.** Broadleaved hedgerow. **D.** Broadleaved forest.

### 2.2 Small mammal survey

Small mammals (i.e. small rodents and shrews) were trapped in spring and autumn of 2023 and 2024, as well as spring 2025. Broadleaved forest sites were included only from spring 2024 onward.

At each site, 20 INRA live traps (160 × 50 × 50-mm aluminium tunnel coupled with a 150 × 70 × 70-mm plastic rest-box) were used. The traps were placed 5 m apart to form a 100 m long line. Coordinates of traps were systematically recorded. Traps were baited with a mixture of sunflower seeds, carrots, and sardines, while the rest-box was filled with cotton to ensure appropriate shelter. Traps were set for three to four consecutive nights and checked every morning during each trapping session.

On the day of capture, rodents were anesthetized with isoflurane and euthanized by cervical dislocation, as recommended by Mills *et al*. (1995). Specimens were weighed and measured (body and tail length), and both external and internal sexual characteristics were recorded. In accordance with our permit conditions, rodents captured beyond the authorized removal number were released following ethical guidelines and marked to avoid recapture bias. Shrews were always released, as they were not targeted for tissue sampling, but systematic marking was only implemented starting in 2024.

All captured individuals were identified to species level using morphological criteria in the field or using molecular methods when necessary (DNA fingerprinting and microsatellite for *Apodemus* species or COI barcoding for other species).

The CBGP laboratory has approval (F-34-169-001) from the Departmental Direction of Population Protection (DDPP, Hérault, France) for the sampling of small mammals and the storage and use of their tissues. All procedures related to small mammals captured in this study complied with the ethical standards of the relevant national and European regulations on the protection of animals used for scientific purposes (Directive 2010/63/EC revising Directive 86/609/EEC). All procedures have undergone validation by the regional ethics committee “Comité d’Ethique pour l’Expérimentation Animale Languedoc Roussillon n°36” in 2023 (num: 2073).

### 2.4 Statistical analyses

Analyses were restricted to the first three trapping nights to standardize effort. Recaptures and empty triggered traps were not considered. All data exploration and statistical tests were performed with R (v4.4.1).

### Habitat types and temporal effects on species richness and relative abundance

#### Species richness

We assessed the effects of habitat type (pine edges, hedgerows or broadleaved forests), season (spring or autumn) and year (2023, 2024 or 2025) on small mammal species richness at the site-session scale. Species richness was modeled using a generalized linear mixed-effects model (GLMM) with a Conway-Maxwell-Poisson distribution. The full model included habitat type, season and year, as well as their interactions with habitat type, and included site identity as a random intercept.

Model selection followed an information theoretic approach using the corrected Akaike Information Criterion (AICc). All submodels derived from the full model were fitted and ranked using the *dredge* function from the *MuMIn* package (v1.48.11). Candidate models were defined as those within two AICc units of the top model. The most parsimonious model among these was retained for inference. Post-fit comparisons were conducted on response-scaled estimated marginal means using Tukey’s range test (α = 0.05).

Models were fitted using *glmmTMB* (v1.1.11), with diagnostics performed using *DHARMA* (v0.4.7) and marginal means estimated with *emmeans* package (v1.11.1).

#### Dominant species relative abundance

Using the same modelling framework, we modeled the relative abundance of the most abundant species, the wood mouse *Apodemus sylvaticus* and the greater white-toothed shrew *Crocidura russula*. Relative abundance was defined as the number of individuals captured per site-session, modeled using a GLMM with a negative binomial distribution. Recaptures were excluded. For unmarked shrews of 2023, the minimum number of individuals captured per line was used (maximum caught in a single night). Scaled spatial coordinates were included as fixed covariates in all submodels to account for large-scale spatial structure.

#### Multi-scale environmental drivers of *A. sylvaticus* abundance in hedgerows

Hedgerow-specific analyses focused on *A. sylvaticus*, the sole dominant species in this habitat type, as *C. russula* was detected in only 26 of 116 site-sessions and at low abundance.

#### Environmental descriptors

We used a multi-scale approach to identify environmental drivers shaping its abundance. We evaluated the relative contribution of climatic conditions, surrounding habitat amount, and local hedgerow characteristics including geometry, composition and structural complexity.

Climatic conditions were assessed using temperature and total precipitation extracted from ERA5 hourly data (0.25° resolution). For each sampling site and session, these climatic variables were aggregated over the 180 days preceding trapping, yielding cumulative precipitation and mean daily temperature as climatic descriptors.

Broadleaved cover was used as a proxy for overall forest habitat in the surrounding landscape of each hedgerow. Using 2022 infrared orthophotographs (20 cm resolution; IGN), we quantified the influence of this land cover around each hedgerow using a Gaussian spatial influence function modelling the effect of vegetation cover on *A. sylvaticus* abundance as a function of distance (Caumette *et al*., 2024) with the *scalescape* package (v0.0.0.9; Lowe *et al*., 2022).

Hedgerow geometry was characterized from 2022 infrared orthophotographs as total length and mean width.

The composition of hedgerow vascular plant communities was surveyed in June 2023 using a standardized protocol described in (Plat *et al*., 2026). Vegetation was recorded by vertical strata and classified as tree (> 5 m), shrub (1-5 m) and herbaceous layers (< 1 m). Species richness of each stratum was considered.

Habitat structural complexity was assessed in 2023 using Airborne Laser Scanning (ALS) data derived from a drone-mounted LiDAR sensor, combined with field measurements of vegetation cover following a protocol described in Plat, Pellet, *et al*. (under review). We considered horizontal cover and its associated variability for the tree and shrub strata using these LiDAR data. Herbaceous cover was estimated using field measurements. Further details regarding environmental descriptors acquisition are provided in Appendix S1.

#### Influence of environmental descriptors on A. sylvaticus relative abundance

As seasonal responses were expected to differ, spring and autumn datasets were analysed separately. Relative abundance was modelled using negative binomial GLMMs. The full model included all environmental descriptors and year as fixed effects. Prior to modelling, multicollinearity was assessed using variance inflation factors (VIF). Variables were sequentially removed until all VIF were below five. All submodels derived from the full model with ΔAICc ≤ 2 were retained as candidates, and multimodel inference was performed using full model averaging based on the AICc and the Akaike weight, which represent the relative likelihood of each model within the candidate set (Burnham and Anderson, 2004).

## 3 RESULTS

### 3.1 Trapping results

Sampling effort comprised 35 sites in spring 2023, 36 in autumn 2023, 52 in spring 2024, and 51 in both autumn 2024 and spring 2025.

Overall, 12,538 effective trap-nights yielded at least 653 distinct individuals, excluding marked individuals and nine potential recaptures among the 37 shrews caught in autumn 2023.

The small mammal community was dominated by two generalist species: *Apodemus sylvaticus* (n = 470, 72% of individuals, detected at 98% of sites) and *Crocidura russula* (n = 133, 20% of individuals, detected at 70% of sites). Other species were much less abundant, with 39 *Clethrionomys glareolus* (bank vole), a forest specialist (6% of individuals, detected at 23% of sites), two *Sorex coronatus*, and single individuals of *Micromys minutus* and *Mus musculus*. Seven shrews could not be confidently identified using molecular methods. Given the low shrew diversity observed among our captures, these individuals were assigned to *C. russula*. *A posteriori* verification confirmed that this assignment had no impact on analyses.

Sampling effort was 3,432 trap-nights in pine edges, 6,435 in hedgerows, and 2,671 in broadleaved forests. Both *A. sylvaticus* and *C. russula* were frequently detected across the three habitat types, whereas the forest specialist *C. glareolus* was almost absent from pine edges and hedgerows, with only three individuals found in these habitats (Figure 2A). *C. glareolus* detection was also strongly seasonal, with 36 of the 39 individuals captured in spring. The single *M. minutus* was captured in a broadleaved forest, the single *M. musculus* in a hedgerow, and the two *S. coronatus* in a hedgerow and a broadleaved forest.

**FIGURE 2.**
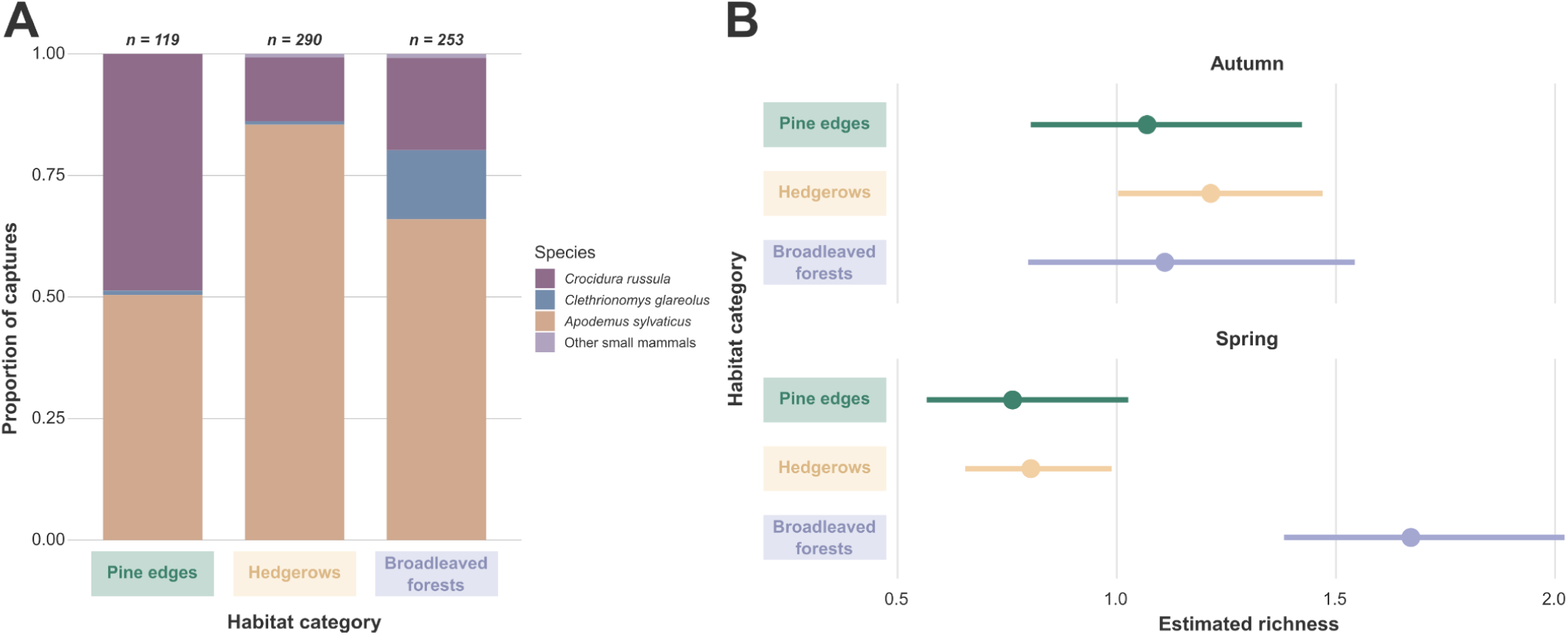
Species composition of small mammal communities by habitat type. The category “Other small mammals” includes the four individuals identified as *Micromys minutus*, *Mus musculus* and *Sorex coronatus*. **B.** Estimated marginal means of species richness per site-session across habitats and seasons. Bars represent 95% CI.

### 3.2 Habitat and seasonal-driven variation

#### Impact on species richness

Local species richness was low, with a mean of 1.08 species per site-session and a range from 0 to 4. The most parsimonious model (Table S1) indicates that small mammal richness varied among habitat types in interaction with season (GLMM_1_; AICc = 510; *df* = 8). In autumn, species richness did not differ among habitat types (all *p* > 0.74; Figure 2B). In contrast, in spring, broadleaved forests supported more than twice the small mammal richness of pine edges (Tukey-adjusted response-scaled contrast: RR = 2.19; 95% CI = [1.52, 3.16]; *p* < 0.001), whereas richness did not differ between hedgerows and pine edges (RR = 1.05; CI = [0.69, 1.62]; *p* = 0.95). Seasonal trends also differed among habitats. Small mammal richness in broadleaved forests was lower in autumn than in spring (RR = 0.66; CI = [0.46, 0.95]; *p* = 0.03), whereas the opposite trend was detected in hedgerows (RR = 1.51; CI = [1.16, 1.96]; *p* = 0.002). Pine edges showed a similar tendency to the latter, though non-significant (RR = 1.40; CI = [0.95, 2.10]; *p* = 0.08).

#### Impact on the abundance of dominant species

The most parsimonious model regarding *A. sylvaticus* shows that its relative abundance varied among habitat types, seasons and sampling years (GLMM_4;_ AICc = 812; *df* = 10). The abundance of *A. sylvaticus* was twice higher in hedgerows than in pine edges (RR = 2.0; CI = [1.25, 3.26]; *p* = 0.0016; Figure 3A) and similar in broadleaved forests (RR = 0.87; CI = [0.54, 1.41]; *p* = 0.78). Contrasts indicated a significant increase of *A. sylvaticus* relative abundance through years (all *p* < 0.01). The species was also more abundant in autumn than in spring (RR = 3.1; CI = [2.17, 4.38]; *p* < 0.0001).

**FIGURE 3.**
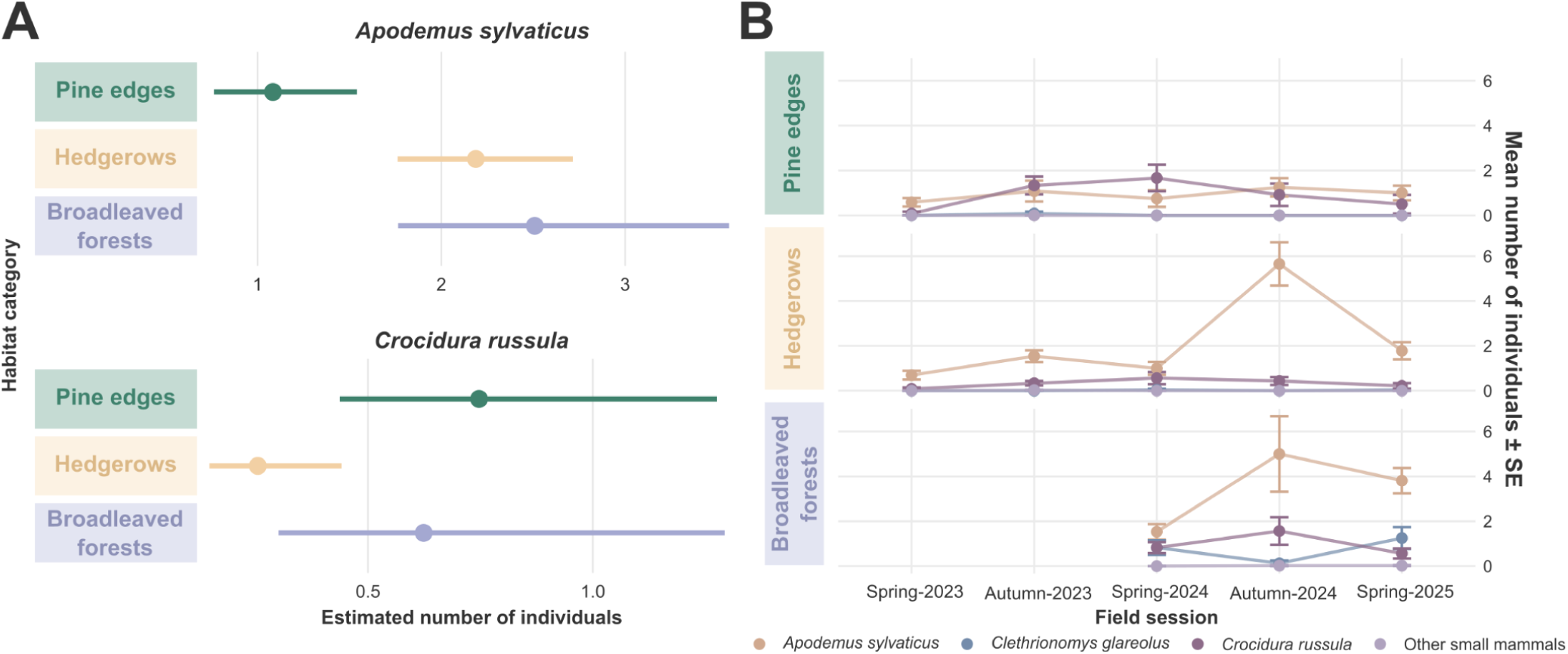
**A.** Estimated marginal means of the individual counts per site-session across habitats and seasons for *Apodemus sylvaticus* (top) and *Crocidura russula* (bottom). Bars represent 95% CI. **B.** Temporal dynamics of the mean number of individuals (± SE) per species and habitat type. The category “Other small mammals” includes the four individuals identified as *Micromys minutus*, *Mus musculus* and *Sorex coronatus*.

Although the interaction between habitat type and year appeared in some competitive models for *A. sylvaticus* (Table S1), it was not retained in the most parsimonious model. Nevertheless, visual patterns strongly suggest that the temporal increase in abundance through years did not occur in pine edges (Figure 3B).

The most parsimonious model regarding *C. russula* indicates that its relative abundance varied among habitat types and years (GLMM_8;_ AICc = 464; *df* = 9). Hedgerows supported lower relative abundance of *C. russula* than pine edges (RR = 0.34; CI = [0.14, 0.79]; *p* = 0.007) and tended to support lower abundance than broadleaved forests, although this trend was not statistically significant (RR = 0.40, CI = [0.15, 1.10]; *p* = 0.087). Relative abundance of *C. russula* peaked in 2024, with a significant decrease detected between 2024 and 2025 (RR = 2.3; CI = [1.10, 5.20]; *p* = 0.045).

### 3.3 Multi-scale environmental drivers of *Apodemus sylvaticus* abundance in hedgerows

Climatic descriptors were strongly correlated with year and with each other in both seasons (VIF > 5). To reduce collinearity, only precipitation was retained in the candidate submodels. All other variables showed Spearman correlation coefficients below 0.6 and were therefore included in the submodels.

Multimodel averaging across candidate models revealed season-specific effects of environmental variables on *A. sylvaticus* abundance (Figure 4 and Table S1). Broadleaved cover influence, width, shrub species richness, and herb species richness were not retained in any candidate models.

**FIGURE 4.**
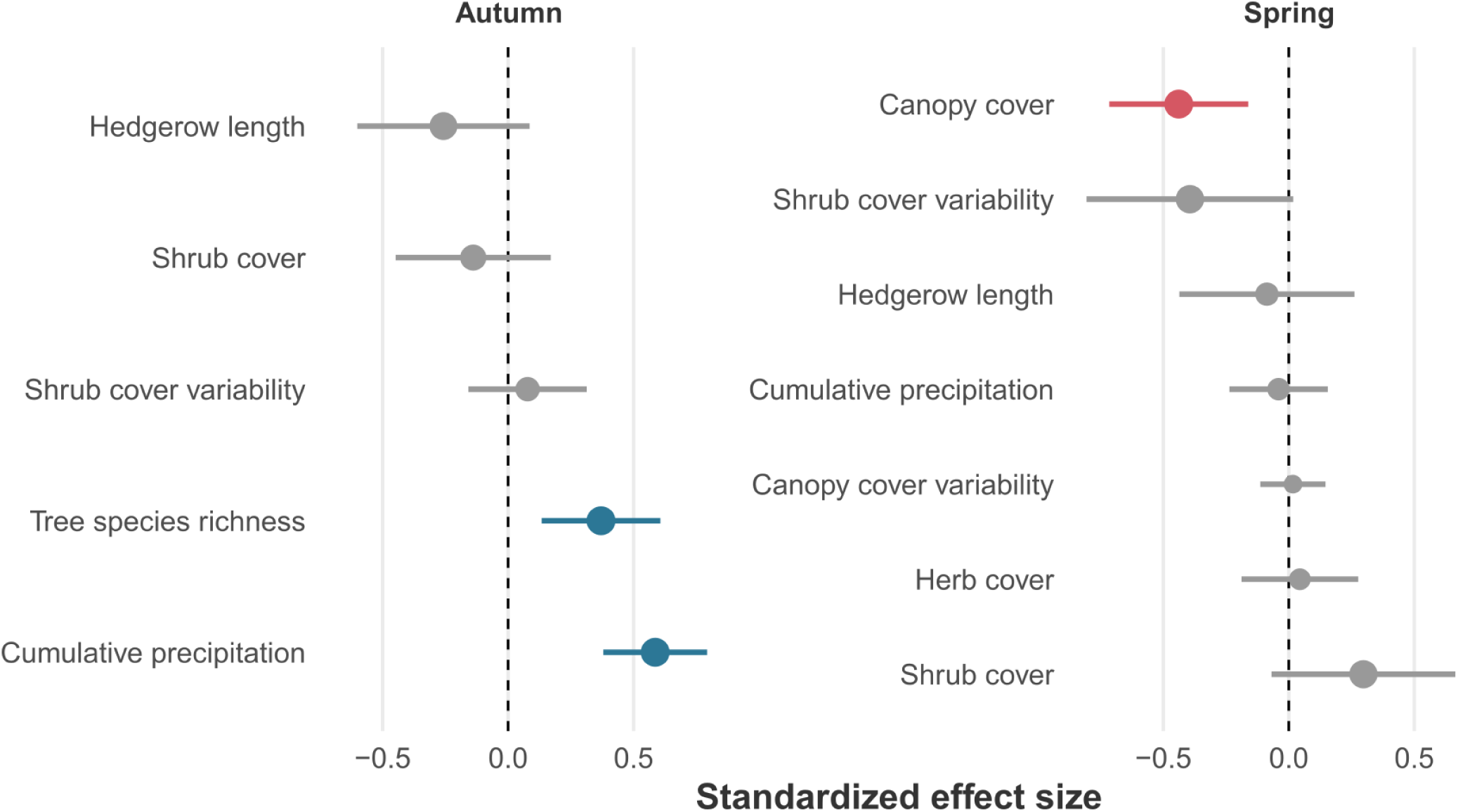
Model averaged standardized regression coefficients (± 95% CI) for the effects of multi-scale hedgerow characteristics on *Apodemus sylvaticus* abundance across seasons. Estimates were derived from candidate models with ΔAICc ≤ 2. Point size indicates the relative importance of each predictor across candidate model sets.

In autumn, abundance increased with cumulative precipitation (β = 0.58; CI = [0.38, 0.79]; *p* < 0.001) and with tree species richness (β = 0.37; CI = [0.13, 0.61]; *p* = 0.002).

In spring, abundance tended to increase with shrub cover (β = 0.29; CI = [-0.07, 0.66]; *p* = 0.11) but decreased with increasing canopy cover (β = -0.44; CI = [-0.71, -0.16]; *p* = 0.002). Additionally, there was weak evidence for a negative relationship with shrub cover variability (β = -0.39; CI = [-0.81, 0.02]; *p* = 0.06).

## 4 DISCUSSION

In this study, we compared small mammal communities across three habitat types within a native pine plantation-dominated landscape, namely pine plantation edges, broadleaved hedgerows and natural broadleaved forests, to assess whether hedgerows could act as biodiversity restoration elements and to determine the characteristics that enhance their effectiveness.

Small mammal species richness was lower in plantations than in natural forest, and the hedgerows studied did not offset this difference. Generalist species dominated these communities in plantations and hedgerows, reflecting the limited suitability of these habitats for forest specialists. However, hedgerows altered community structure by modifying the relative abundance of dominant species, with responses varying among taxa.

Community patterns were further shaped by rodent demographic dynamics. Overall, our findings highlight both the potential and limitations of hedgerows as restoration elements in plantation landscapes, and emphasise the need to consider species-specific responses and temporal variability when designing, evaluating and monitoring biodiversity-oriented management interventions.

### 4.1 Hedgerows do not increase small mammal richness in pine plantations

Sampling across 52 sites over two and a half years yielded six different species of small mammals within the study area. Communities were over-dominated by two generalist species : the wood mouse (*Apodemus sylvaticus*) and the greater white-toothed shrew (*Crocidura russula*), which accounted for more than 90% of captured individuals and were consistently detected across sites. Such dominance across sites suggests strong environmental filtering favouring disturbance-tolerant species with broad niches, flexible diets or sufficient dispersal capacity.

Pine plantation edges and broadleaved hedgerows adjacent to plantations supported lower species richness than natural broadleaved forests. This pattern was mainly driven by the near absence of the forest specialist bank vole (*Clethrionomys glareolus*), resulting in communities solely composed of generalists. This species is not inherently excluded from conifer forest and can occur in pine stands under suitable microhabitat conditions (Birkan, 1968). Rather than tree species identity *per se*, dense understorey appears to be the key determinant of their habitat suitability (Appleby and Balkenhol, 2024).

*Clethrionomys glareolus* is typically associated with dense and structurally complex understories that provide both trophic resources and protective cover (Mazurkiewicz, 1994). Although this species exhibits relatively broad trophic breadth, it relies heavily on the consumption of green plant material from the herbaceous layer (Butet and Delettre, 2011). Young pine stands with well-developed ground vegetation can sustain higher bank vole abundance than older stands with sparse understorey (Birkan, 1968). In our maritime pine plantation sites, the simplified understorey due to high tree density and mechanical clearing likely contributes to the exclusion of this forest-associated specialist from the regional pool.

In contrast, *A. sylvaticus*, is more granivorous and exhibits strong dietary plasticity, enabling it to exploit a wide range of resources across multiple vegetation strata (Butet and Delettre, 2011). Such ecological flexibility allows the species to adjust its diet to local resource availability, exploit multiple habitat types (Unnsteinsdottir and Hersteinsson, 2011), and persist within low-quality matrices that other forest species avoid (Heroldová *et al*., 2007). For example, its increased consumption of animal prey in coniferous habitats has been interpreted as a compensatory response to reduced fruit availability (Montgomery and Montgomery, 1990). Such trophic flexibility likely contributes to its presence in conifer plantations (Gasperini *et al*., 2016) despite ecological simplification.

Despite being relatively unmanaged, more diverse, and structurally complex than plantation edges (Plat *et al*., 2026) the broadleaved hedgerows in our study area supported almost no *Clethrionomys glareolus*, even though such linear habitats can constitute the sole habitat for this species in other productive landscapes (Heroldová *et al*., 2007). This likely reflects a combination of remaining ecological constraints that outweighed hedgerows’ potential benefits. Increased plant diversity does not always translate into suitable trophic resources. Moreover, as permanent edge habitats, hedgerows face reduced microclimatic buffering compared to forest interiors (Vanneste *et al*., 2020), potentially maintaining suboptimal conditions for forest specialists. However, local characteristics alone are unlikely to explain the observed pattern. *C. glareolus* has been reported to be more abundant at forest edges adjoining large woodlands than in isolated hedgerows (Schlinkert *et al*., 2016) suggesting that habitat configuration and landscape context are critical determinants of species occurrence. The narrow configuration of hedgerows may prevent them from sustaining viable populations, as small patches are more exposed to demographic stochasticity and local extinctions. Indeed hedgerows have sometimes been described as lower-quality woodland habitats (Hinsley and Bellamy, 2000) or demographic sinks (Guivier *et al*., 2011). In this context, population persistence in hedgerows may depend on recolonisation from nearby source habitats.

Landscape constraints likely further limited the suitability of hedgerows as habitats for disturbance-sensitive species. Broadleaved and mixed stands represented less than 15% of forest cover in the study area, and within a 500 m radius around hedgerows, accounted for only 10% of the area on average, potentially limiting the availability of source populations. Rare and small broadleaved stands within the 890 km² plantation matrix are likely insufficient to sustain stable *C. glareolus* populations, with only larger patches functioning as persistent sources. In human-modified landscapes, forest-dwelling species are often reported to decline sharply below 30-40% forest cover (Arroyo-Rodríguez *et al*., 2020), suggesting that limited extent of broadleaved cover may have constrained the occurrence of *C. glareolus*, particularly within hedgerows.

### 4.2 Hedgerows shift dominant species composition

We revealed that hedgerows transformed community composition by their impact on relative abundance of generalist species. Shrews were less frequently captured in hedgerows than in pine edges, whereas *A. sylvaticus* reached relative abundances comparable to those observed in broadleaved forests.

Presence alone does not necessarily indicate that a habitat is suitable or optimal. Although *A. sylvaticus* can occur in conifer-dominated forests, its abundance is frequently lower than in broadleaved or mixed forests (Navarro-Castilla and Barja, 2019; Appleby and Balkenhol, 2024). While this species benefits from understorey complexity, its population density is more tightly linked to tree species composition and associated seed resources than *C. glareolus* (Appleby and Balkenhol, 2024). A*podemus* spp. rely heavily on large, energy-rich seeds, particularly acorns, which constitute a substantial proportion of their diet in broadleaved systems (Watts, 1968). Although they occasionally consume conifer seeds, including pinecones (Manso *et al.,* 2014), seed biomass is generally lower in pine-dominated stands than in oak-dominated ones (Navarro-Castilla and Barja, 2019), and conifer seeds may contain secondary compounds that reduce palatability or nutritional value (Lobo, 2014). Reduced seed quantity and quality may therefore lower carrying capacity in pine plantations. Understorey plant species richness is also higher in hedgerows than in pine edges (Plat *et al*., 2026), potentially increasing food resource diversity. Overall, the higher abundance of *A. sylvaticus* in broadleaved forests and hedgerows than pine edges likely reflect the combined effect of greater tree seed production and more complex understorey. Hedgerows may further function as refuge habitats following disturbances in adjacent pine stands, as described in agricultural landscapes (Tattersall *et al*., 2001).

In contrast, *C. russula* exhibited lower relative abundance in hedgerows than in plantations and broadleaved forests, although variations were more modest. As a thermophilous insectivore associated with warm and open habitats that provide sufficient ground cover for protection (Torre *et al*., 2020), the species did not appear constrained by plantation environments in our study area. Pine plantations may offer favourable microclimatic conditions at ground level, while maintaining sufficient herbaceous cover and invertebrate availability. Hedgerows, with their denser canopy and their cooler microclimatic conditions, may be less suitable for this thermophilous species. Invertebrate communities also differ from those at pine edges (e.g. ground-dwelling spiders; Plat *et al*., 2026), potentially resulting in different prey assemblages. Although broadleaved forests and hedgerows share comparable tree composition, the sampled forests included heterogeneous environments, with canopy gaps and semi-open patches that likely created suitable microhabitats. This heterogeneity may explain why *C. russula* maintained comparable abundance in plantations and forests but not in hedgerows. Given the limited ecological information currently available for this species, longitudinal survey and mechanistic studies are needed to clarify how thermal conditions, vegetation structure and prey availability interact to shape *C. russula* abundance and distribution in this landscape.

Altogether, the shifts in relative abundance of the dominant species suggest that hedgerows may support greater overall small mammal biomass than plantation stands. Although they did not increase species richness, their effect on abundances may nonetheless have ecological consequences as increased prey biomass can positively affect predator populations (Šálek *et al*., 2010) or epidemiological dynamics (Keesing and Ostfeld, 2021).

### 4.3 Seasonal and interannual population dynamics shape biodiversity patterns within and between habitats

Short-lived rodents are well-known for exhibiting marked temporal fluctuations in population dynamics. In seasonal environments, populations typically show strong intra-annual fluctuations driven by changes in reproduction and survival. Seasonal dynamics can overlap with long-term interannual dynamics, including episodic outbreaks and, in some species, multiannual population cycles (Andreassen *et al*., 2021).

In our study, seasonal variation in species richness across habitats partially reflected these dynamics. Two distinct patterns emerged. First, species richness within habitats showed seasonal fluctuations, in pine edges and hedgerows only. The higher richness detected in autumn was driven by the growth and wider occurrence of *A. sylvaticus* populations across sites. This pattern aligns with well-documented seasonal dynamics in temperate northern European populations, where reproduction from spring to summer leads to an autumn peak in abundance, followed by a winter decline (Bergstedt, 1965).

Second, differences in species richness among habitats were detectable only in spring, when *A. sylvaticus* occurred at lower frequencies and *C. glareolus* predominated in captures. The spring predominance of *C. glareolus* diverges from the patterns commonly observed in northern populations and may indicate shifts in reproductive phenology or overwinter survival. Reproductive seasonality in rodents is highly plastic, potentially varying with latitude, climatic conditions and resource availability (Heldstab, 2021). In its southernmost range, contrasted seasonal dynamics have been reported for *C. glareolus*, with breeding concentrated from autumn to spring in Mediterranean systems (Huerta-Schliemann *et al*., 2025). Such plasticity, combined with documented differences in reproductive seasonality among sympatric rodents (Massoud *et al*., 2021), suggests that environmental conditions may alter the seasonal contribution of each species to community composition.

Beyond seasonal variation, we observed yearly variations in the relative abundance of *A. sylvaticus*. Wood mice are prone to irregular interannual outbreaks, often associated with pulsed resource availability, particularly mast seeding events (Selås, 2020). As plant productivity and masting are shaped by preceding climatic conditions (Fleurot *et al*., 2023), these fluctuations can generate strong bottom-up effects, resulting in marked year-to-year variation in rodent abundance (Ferrari *et al*., 2025). In our study, the population peak appeared mainly in broadleaved hedgerows, but not in pine edges. Although the interaction between year and habitat type was not retained after model selection, this pattern is consistent with the strong dependence of *Apodemus* spp. on tree seeds. While the link between rodent demography and masting is well established in broadleaved systems, evidence for consistent responses to conifer masting is limited (Lobo, 2014), suggesting that pine-dominated environments may not provide the resource pulses that influence seed-dependent rodents. Interannual population variation may also reflect direct influence of climatic extremes on survival and reproduction (Huerta-Schliemann *et al*., 2025). Severe droughts, heatwaves, and large wildfires affected western Europe and the Landes forest in 2022-2023 (Toreti *et al*., 2022; Menut *et al*., 2023). Given their weaker microclimatic buffering, pine plantations may exacerbate the consequences of such events, thus contributing to the lower abundances observed in these environments.

Overall, our results highlight that habitat suitability operates within a temporally dynamic framework where reproduction, resource pulses and climatic extremes interact to shape communities across years and seasons. Therefore, habitat effects on small mammals cannot be fully understood without accounting for demographic processes and monitoring communities over time. Snapshot surveys may fail to detect habitat-dependent responses; for instance, had sampling been restricted to 2023, differences in *A. sylvaticus* abundance among habitat types would not have been detected.

### 4.4 Hedgerows have limited effects on biodiversity recovery in pine monoculture

By comparing small mammal communities across broadleaved habitats within the monocultural Landes forest, our results provide insight into the potential of hedgerows as a nature-based solution to mitigate biodiversity loss in pine plantation-dominated landscapes.

Our results suggest that hedgerows alone may not substantially increase small mammal species richness in pine plantations, but can provide complementary habitat or refuges for species poorly represented in the plantation matrix. In practice, integrating hedgerows into plantation-dominated landscapes could enhance connectivity and support source populations. Our results testify that the ability of hedgerows to promote biodiversity depends on both local habitat quality and landscape context. The case of *Apodemus sylvaticus* illustrates how hedgerows’ attributes can shape the ecological responses of small mammals throughout their annual cycle. In autumn, abundance increased with precipitation and tree species richness, consistent with bottom-up effects mediated by resource availability through enhanced primary productivity and seed resource diversity. In spring, abundance was primarily associated with vegetation structure: well developed and continuous shrub layers were favoured, whereas dense canopy cover was detrimental (e.g., Jaime-González *et al*., 2017). A structured understorey likely enhances shelter and access to early vegetative resources during a period when seeds are scarce and survival constraints are strong. These patterns suggest that hedgerows’ resources may limit populations during the reproductive phase, whereas their structural cover becomes more important for overwintering and early-spring survival. Altogether, this implies that hedgerows design and management are essential determinants of their ecological value. Optimising establishment attributes (e.g. width, tree composition) or applying management practices that enhance their quality as habitats (e.g. coppicing to promote dense shrub layers) could improve suitability for a broader range of species (Graham *et al*., 2018).

Here, several factors likely limited the potential of hedgerows to enhance small mammal biodiversity in our study. First, the relatively low species diversity in the regional pool (Ruys and Couzi, 2015) may have limited the detection of hedgerow benefits as higher regional biodiversity increases the likelihood that functionally distinct species occupy the niches provided by intermediate-quality habitats. Second, the hedgerows we sampled were neither established nor managed for conservation purposes. As such, their attributes and landscape configuration may not have provided the optimal conditions for forest specialists. Third, most of the sampled hedgerows were isolated, with few other broadleaved elements embedded within the pine plantation matrix. Local optimisation alone may therefore be insufficient to overcome strong landscape constraints. Increasing hedgerow density to approach a “bocage” landscape (Boinot *et al*., 2023), or restoring larger broadleaved patches, could enhance habitat amount, connectivity and environmental heterogeneity (Forman and Baudry, 1984), all of which are key drivers of biodiversity in fragmented landscapes. Finally, given the scarcity of broadleaved elements in the plantation landscape, sporadic hedgerow establishment is unlikely to increase small mammal species richness. Coordinated, landscape-scale planning and sustained incentives for hedgerow deployment may therefore be necessary to move beyond small-scale habitat enhancement toward functional habitat networks.

Overall, our study highlights that the effectiveness of hedgerow integration as a nature-based solution for maintaining or restoring biodiversity cannot be assessed solely on the basis of small mammal species richness. Changes in abundance patterns may precede shifts in species richness and can provide early indicators of ecological response to interventions, particularly in ecologically simplified systems such as the pine monoculture of the Landes forest, France. Monitoring efforts should therefore extend beyond diversity metrics to include demographic responses and population dynamics. Temporal surveys spanning multiple years are essential, as community composition and demographic processes may take time to reach equilibrium following habitat modification (Jackson and Sax, 2010), and interannual variability can strongly affect species occurrence and abundance. Finally, evaluating responses across multiple trophic levels may help capture cascading effects and provide a more comprehensive understanding of the ecological impacts of nature-based solutions such as habitat restoration.

## Author contributions

Aloïs Berard, Hervé Jactel, Jean-Baptiste Rivoal, Nattan Plat and Nathalie Charbonnel conceived the ideas and designed methodology. Aloïs Berard, Anne Loiseau, Cameron Pellett, Jean-Baptiste Rivoal, Julien Pradel, Julie Blanchet, Karine Berthier, Lily Cesari, Maxime Galan, Nathalie Charbonnel, Nattan Plat, Ruben Valbuena and Sylvain Piry contributed to data collection and curation. Aloïs Berard performed the analyses and led the writing. Nathalie Charbonnel supervised the project and acquired the funding. All authors contributed to manuscript improvements and gave final approval for publication.

Whenever possible, results were discussed with local stakeholders to enhance involvement in the co-design of future interventions and share the naturalist data collected.

## Acknowledgements

We are grateful to the Réserve Géologique de Saucats et La Brède National Nature Reserve management team and the National Forests Office for their support in granting permissions and offering guidance. We would also like to thank Florent Sebbane and the Pasteur-Lille team for their support during fieldwork. This project has received funding from the European Union (grant agreement No 101060568; project BEPREP, grant agreement No 101036849; project SUPERB, grant agreement No 101059498; project eco2adapt).

## Data availability

The occurrence dataset, including individual characteristics, is available on GBIF (Berard *et al*., 2026). The associated biological samples have been incorporated into the CBGP reference collection of small mammals (CBGP, 2025). The scripts and datasets used for the analyses are available on Zenodo (DOI to be provided upon acceptance).

