## Appendix S1 for "Broadleaved hedgerows as complementary habitats for small mammals in pine plantation landscapes"

### Appendix S1: Data acquisition and processing of multi-scale environmental drivers in hedgerows

#### 1. Surrounding landscape

High-resolution infrared colour orthophotographs (IGN; 20 cm pixel resolution, https://geoservices.ign.fr/documentation/donnees/ortho/bdortho) were analysed at a scale of 1:2500 in a GIS environment to map broadleaved and mixed forest stands surrounding each hedgerow.

To quantify the influence of surrounding broadleaved forest on Apodemus sylvaticus abundance, we modelled the distance decay of broadleaved forest cover using a Gaussian spatial influence function. The optimal spatial scale was estimated using a distance weighted modelling framework implemented in scalescape package (with a maximum radius of 1 km). For this, abundance variation was modelled using a negative binomial distribution with year and season included as covariates. Negative binomial models were fitted using the MASS package.

Using the estimated Gaussian function, we then calculated the cumulative broadleaved forest cover effect for each hedgerow by applying the distance based weights to surrounding broadleaved forest cover and summing the weighted values.

#### 2. Geometry

Hedgerow geometry was quantified using total length and mean width. For each hedgerow, a GIS-derived polygon was created at 10 m resolution from 20 cm resolution 2022 infrared orthophotographs (IGN). The shapefile followed the crown outline of each hedgerow.

Width was estimated along the same 100 m section of polygon where biodiversity surveys were conducted. A centreline representing the longitudinal axis of each hedgerow was derived, and a dense set of segments perpendicular to this axis was generated. Only segment that did not intersect with many other segments were retained. The length of the remaining segments was measured, and the mean of these values was used as the width estimate for each hedgerow.

#### 3. Botanical composition

Vascular plant communities were surveyed in June 2023 in each hedgerow using three 5 × 1.5 m quadrats evenly spaced along a 100 m section.

Vegetation was classified into three vertical strata:

- Herbaceous layer: herbaceous and ligneous species under 0.5 m in height
- Shrub layer : only ligneous species and climbers (vines) between 0.5 and 5 m in height
- Tree layer only ligneous species and climbers (vines) above 5 m in height.

Species were identified in the field by two pairs of trained observers using a taxonomic guide (Flore forestière française; Rameau et al., 2018). When identification was uncertain, specimens were collected and verified in the laboratory using a binocular microscope.

#### 4. Structural complexity

Airborne Laser Scanning (ALS) data were acquired in summer 2023 using a drone-mounted LiDAR sensor (DJI Matrice 300 RTK with a zenmuse L1 LiDAR sensor). Twenty two of the 24 hedgerows were fully scanned.

For each hedgerow, the LiDAR point cloud was clipped to the same 100 m polygon used for width estimation. Raw point clouds were processed using the Point Cloud Data Abstraction Library (PDAL) as follows. Ground points were classified using a simple morphological filter (smrf; Pingel et al., 2013). The inverse distance weighted average of ground point heights was used to create a 2 m resolution digital terrain model (dtm). All points within the hedgerow polygon could then be height normalised to distance above the dtm. Finally points with a greater scan angle than 20 degrees were removed

To characterise spatial variation in vegetation structure, Voronoi tessellation was applied along the hedgerow centreline. Polygon centres were spaced at least 10 m apart, resulting in a median of eight polygons per hedgerow. In some cases, up to two polygons more or fewer were generated depending on hedgerow shape and width.

Within each polygon, the vegetation cover was estimated from the point cloud for canopy (5m to maximum height) and shrub cover (1-5m). For each hedgerow, mean cover and along hedgerow variability were calculated from polygon level estimates (Plat et al., submitted).

Herbaceous cover was quantified using ground-based measurements rather than LiDAR, as near ground returns can be biased. Herbaceous cover was estimated as the vertical projection of the herbaceous stratum using a Domin scale assessment conducted during botanical surveys (three quadrats along 100 m). Variability in herbaceous cover along hedgerows was not considered as the number of sampled quadrats was considered too low to accurately capture variability (Plat et al, submitted).
